# Kappa-opioid receptor-dependent changes in dopamine and affective behavior occur differentially across the nucleus accumbens shell rostro-caudal axis

**DOI:** 10.1101/2020.05.09.085761

**Authors:** Breanne E. Pirino, Mary B. Spodnick, Andrew T. Gargiulo, Genevieve R. Curtis, Jessica R. Barson, Anushree N. Karkhanis

**Author notes:** Corresponding author. Department of Neurobiology and Anatomy, Drexel University College of Medicine, Philadelphia, PA 19129, USA.,E-mail address (J.R. Barson). Corresponding author. Department of Psychology, Binghamton University – SUNY, Binghamton, NY 13902, USA.,E-mail address (A.N. Karkhanis). (BEP), (ATG), (GRC), (MBP).

## Abstract

Neural circuit engagement within the nucleus accumbens (NAc) shell is implicated in the regulation of both negative and positive affect. Classically, the dynorphin/kappa opioid receptor (KOR) system in the NAc was believed to promote dysphoric behavior, while dopamine was viewed as interacting with reward behavior, and KOR activation was known to inhibit dopamine release. Recently, however, both the KOR and dopamine systems have, separately, been shown to have differential effects across the rostro-caudal axis of the NAc shell on hedonic responses. Whether or not this is due to interactions between KORs and dopamine, and if it extends to other affective behaviors, remains to be determined. In this study, we examined in rats the relationship between the KOR and dopamine systems in both the rostral and caudal NAc shell using *ex vivo* fast scan cyclic voltammetry and the impact of KOR activation on affective behavior using approach-avoidance assays. We report here that activation of KORs in the caudal NAc shell significantly inhibits dopamine release, stimulates novelty-induced rearing behavior, increases avoidance behavior, and reduces locomotor activity. In contrast, activation of KORs in the rostral NAc shell inhibits dopamine release to a lesser extent and instead increases approach behavior. Taken together, these results indicate that there is heterogeneity across the rostro-caudal axis of the NAc shell in the effects of KOR stimulation on affective behaviors, and they suggest that this might be due to differences in KOR control over dopamine release.

## 1. Introduction

The nucleus accumbens (NAc) processes responses to both aversive and rewarding stimuli, but the mechanisms through which this occurs remain to be determined. A heterogenous area, the NAc receives dopaminergic inputs from the ventral tegmental area, and contains medium spiny neurons and local collateral axons that release dynorphin [1], which acts primarily at the kappa opioid receptor (KOR) [2]. Mesolimbic dopamine, dynorphin/KORs, and their interplay are involved in affective states [3,4]. Dopamine release in the NAc is facilitated by both reinforcing and aversive stimuli [5], and KOR activation promotes both preference and aversion [6]. Thus, both dopamine and the KORs appear to be involved in the ability of the NAc to process aversive and rewarding stimuli, but how this occurs remains to be determined.

Activation of KORs in the NAc shell has only recently been found to have opposing effects on affective behavior [7,8]. In the middle NAc shell of mice, optical stimulation of dynorphin-releasing medium spiny neurons in the ventral subregion was found to promote real-time place aversion, while stimulation in the dorsal subregion was found to augment place preference [7]. In rats, KOR activation in the caudal half of the NAc shell suppressed hedonic orofacial reactions to sucrose while activation in the rostral half enhanced them [8]. These studies suggest that the distinct contributions of KOR activation to affective behavior occur via different anatomical subregions of the NAc shell, but whether this extends to other approach-avoidance behaviors, and what the neurochemical mechanism behind this may be, remains to be determined.

Within the NAc, KORs are known to regulate dopamine. Acting as inhibitory G-protein coupled receptors [9,10], KORs are located on dopamine terminals [11] and their activation inhibits dopamine release [4,12–15]. While dopamine in the NAc is responsive to reinforcing stimuli [16–18], it has recently also been found to be responsive to aversive stimuli [5,19,20]. Furthermore, while dopamine release in the rostral NAc shell inhibits taste aversion, dopamine in caudal shell instead enhances it [21]. Thus, like KORs, dopamine is involved in both aversive and rewarding responses, and this appears to occur via different subregions of the NAc shell.

In this study, we determined if the opposing effects of KOR activation in the NAc shell extended to affective behavior in exploration-based tasks and we examined a potential mechanism for these differences. Our hypothesis was that, in the rostral compared to caudal subregion of the NAc shell, KOR activation results in different or even opposite effects on affective behavior and dopamine transmission.

## 2. Material and methods

### 2.1. Subjects

Adult, male and female Long-Evans rats (*N* = 43; 38 males and 5 females; 8 weeks at the start of testing, Charles River Laboratories International, Inc., Malvern, PA, USA and Kingston, NY, USA) were pair housed (for neurobiological studies) or individually housed (for behavioral studies) in AAALAC-accredited facilities on a 12-hour reversed light/dark cycle (lights off at 0300 h for neurobiological studies or 0900 h for behavioral studies). They were given at least one week to acclimate to the facility and were handled daily prior to the start of experiments. Rats received *ad libitum* chow (Laboratory Rodent Diet 5001, Lab Diet, St. Louis, MO, USA) and water. Experiments were approved by the Institutional Animal Care and Use Committees of SUNY Binghamton University and Drexel University College of Medicine and followed the NIH Guide for the Care and Use of Laboratory Animals.

### 2.2. Drugs

The selective KOR agonist, U50,488 [22,23] was acquired from Tocris (Minneapolis, MN, USA). For neurobiological studies, it was dissolved in 18.0 MΩ water and bath-applied at cumulative concentrations of 0.01 − 1.0 µM. For behavioral studies, it was dissolved in 0.9% saline (Baxter International Inc., Deerfield, IL, USA) and microinjected at 8.0 nmol.

### 2.3. Ex vivo fast scan cyclic voltammetry (Experiment 1)

To determine the effects of KOR stimulation on dopamine dynamics in the rostral compared to caudal NAc shell, *ex vivo* fast scan cyclic voltammetry (FSCV) was used to measure dopamine release under baseline conditions and in the presence of U50,488. Briefly, male and female rats (*n* = 5/group) were sacrificed 6 hours into the dark cycle and their brains were rapidly removed and prepared as previously described [24]. A vibrating tissue slicer was used to prepare 400 μm thick coronal brain sections containing NAc. These slices were immersed in oxygenated artificial cerebrospinal fluid (aCSF; 32°C) containing: 126 mM NaCl, 2.5 mM KCl, 1.2 mM NaH_2_PO_4_, 2.4 mM CaCl_2_, 1.2 mM MgCl_2_, 25 mM NaHCO_3_, 11 mM glucose, 0.4 mM L-ascorbic acid and the pH was adjusted to 7.4. Endogenous dopamine release was evoked by single electrical pulse stimulation (monophasic+, 4 ms, 750 µA) applied to the tissue every 5 minutes and dopamine was detected using a triangular scanning waveform (−0.4 to +1.2 and back to -0.4 V, Ag *vs* AgCl) at the rate of 400 V/s. Recording electrodes were calibrated with known concentrations of dopamine (3 μM). After stable dopamine responses were obtained, cumulative concentrations of U50,488 were bath applied to rostral and caudal NAc shell slices. All FSCV data were analyzed using a Michaelis-Menten kinetics-based algorithm [25], using Demon Voltammetry and Analysis software [26].

### 2.4. Microinjections (Experiments 2 – 4)

Rats were cannulated bilaterally for injection in the rostral or caudal NAc shell, using published methods [27]. Stainless steel guide shafts (21−gauge, Small Parts) were implanted perpendicularly, aimed at the rostral or caudal NAc shell (2.1 mm or 1.1 mm anterior to bregma, ±0.8 mm lateral to midline, 4.0 mm ventral to the level skull) [28]. To prevent occlusion, 26−gauge stainless steel stylets were left in the guide shafts between injections. During their one-week recovery period, rats were handled daily, and their stylets were removed and replaced to acclimate them to the microinjection procedure.

Injections of freshly-prepared U50,488 or saline vehicle were given 2 hours into the dark cycle through concentric microinjectors of 26−gauge stainless steel outside and fused-silica tubing inside (74 μm ID, 154 μm OD; Polymicro Technologies, Phoenix, AZ, USA) that protruded 3.0 mm beyond the guide shafts to reach the NAc shell. A syringe pump (Harvard Apparatus, Holliston, MA, USA) delivered 0.3 μl of solution over 30 seconds, and the microinjector remained in place for an additional 30 − 60 seconds to allow for diffusion. Each hemisphere was injected sequentially for each subject. We have previously demonstrated that injection of methylene blue dye at the volume used (0.3 μl) has a radial spread of approximately 0.5 mm [29,30]. Behavioral testing for each subject began 15 minutes following the completion of their second injection.

### 2.5. Novelty-induced locomotor activity (Experiment 2)

To determine the effects of KOR stimulation on novelty-induced locomotor activity, male rats (*N* = 33) were injected in the rostral or caudal NAc shell between-subject with U50,488 or saline (*n* = 6 – 12/group) and novelty-induced locomotor activity was measured in an activity chamber with an area of 43.2 cm x 43.2 cm with 42 cm high walls (Med Associates, Inc., St. Albans, VT, USA), under very low light (< 5 lux). Animals were placed at the center of the novel chamber and allowed to explore for 15 minutes while vertical counts and ambulatory distance were recorded using infrared beams [27].

### 2.6 Light-dark box (Experiment 3)

To determine the effects of KOR stimulation on approach-avoidance behavior, some rats from Experiment 2 (*N* = 17) were, one week later, injected in the rostral or caudal NAc shell within-subject with U50,488 or saline (*n* = 6 − 11/group) and tested in a light-dark box. This box was the same chamber as used for novelty-induced locomotor activity, but with a two-chamber light-dark insert (Med Associates, Inc., St. Albans, VT, USA). The light chamber was illuminated by LED lamp at 400 lux in the center. Animals were acclimated to the light-dark box in one 5-minute session on the day before testing. On test days, animals were placed in the corner of the light chamber, facing away from the dark chamber, and allowed to explore the light-dark box for 5 minutes, while time spent in and number of entries into the light chamber were recorded using infrared beams. Research suggests that behavior in the light-dark box does not significantly change on repeat testing [31,32].

### 2.7. Open field (Experiment 4)

To determine the effects of KOR stimulation on another measure of approach-avoidance behavior, rats (*N* = 20) were injected in the rostral or caudal NAc shell within-subject with U50,488 or saline (*n* = 7 − 13/group) and tested in an open field. While 11 of these rats were tested one week after the novel activity chamber (Experiment 2), 9 were tested one week after the light-dark box (Experiment 3). The open field was an arena of 81 cm x 81 cm with 31.5 cm high walls (Med Associates, Inc., St. Albans, VT, USA). Testing was conducted under ambient room light of 200 lux. Animals were acclimated to the open field in one 10-minute session on the day before testing. On test days, animals were placed at the center of the open field and allowed to explore for 10 minutes, while time in the center area, number of entries into the center, and overall ambulatory distance were recorded using infrared beams. Behavior in an open field does not significantly change on repeat testing [33].

### 2.8. Histological analysis

Rats were anesthetized with isoflurane, sacrificed by rapid decapitation, and their brains extracted. Microscopic examination of 30 µm coronal sections was used to ensure that injections were made into the NAc shell and determine the level at which this occurred. Bregma +2.76 − +1.80 mm was considered to be rostral and bregma +1.68 – 0.72 mm was caudal [28].

### 2.9. Statistical analysis

Statistical analysis of the FSCV data was conducted using GraphPad Prism 8 (GraphPad Software, La Jolla, CA, USA). Baseline dopamine release and uptake data were analyzed using paired two-tailed Student’s *t*-tests, as this was a within-subject design. The U50,488 concentration response curves were analyzed using a repeated measures (RM) two-way analysis of variance (ANOVA). The dependent variable was [DA]/p and the independent variables were rostral/caudal subregions of NAc shell and U50,488 concentration. Significant effects were followed up by Sidak’s pairwise *post-hoc* analysis. Analysis of behavioral data was conducted using IBM SPSS Statistics version 24 (IBM, Armonk, NY, USA). Because the FSCV analyses detected differences between the subregions, an *a priori* decision was made to separately test each subregion. Thus, novelty-induced locomotor activity data were analyzed using unpaired, two-tailed *t*-tests and light-dark box and open field data were analyzed using paired two-tailed *t*-tests. All data are reported as mean ± standard error of the mean. Significance was set at *p* < 0.05.

## 3. Results

### 3.1. Experiment 1: Kappa opioid receptor-mediated inhibition of dopamine release is greater in caudal compared to rostral NAc shell

To determine the effects of KOR stimulation on dopamine dynamics in the rostral compared to caudal NAc shell, dopamine release under baseline conditions and in the presence of U50,488 was measured in female (**Figure 1**) and male (**Figure 2**) rats (*n* = 5/group), using *ex vivo* FSCV. After obtaining stable baseline levels of dopamine release, cumulative concentrations of U50,488 were bath applied to rostral and caudal NAc shell-containing slices. For experiments in females, electrode placements in the NAc shell and representative dopamine signals, along with their respective colorplots, are shown in **Figures 1A** and **1B**, respectively. Comparison of dopamine release between rostral and caudal NAc shell at baseline (pre-drug exposure) revealed no significant differences [*t*(4) = 0.277; *p* = 0.7956] (**Figure 1C**). Similarly, comparison of dopamine uptake rate (Vmax) between the two subregions also did not reveal significant differences [*t*(4) = 0.560; *p* = 0.6055] (**Figure 1D**). On the other hand, for KOR-activation mediated inhibition of dopamine release, a repeated-measures two-way ANOVA comparison of the rostral and caudal subregions of the NAc shell revealed main effects of U50,488 concentration [*F*(4, 32) = 74.890, *p* < 0.0001] and subregion [*F*(1, 8) = 5.960, *p* = 0.0405], as well as a significant interaction between U50,488 concentration and subregion [*F*(4, 32) = 2.930, *p* = 0.0357]. *Post-hoc* Sidak’s pairwise comparisons revealed that U50,488 at 0.3 µM (*p* = 0.0479) and 1.0 µM (*p* = 0.0325) inhibited dopamine release to a significantly greater extent in the caudal compared to rostral NAc shell (**Figure 1E**). The maximal effect of U50,488-induced inhibition of dopamine release was significantly greater in the caudal compared to rostral subregion of the NAc shell [*t*(4) = 8.246; *p* = 0.0012] (**Figure 1F**). A paired *t*-test comparison of the EC50 between rostral and caudal subregions also showed a significant difference [*t*(4) = 7.266, *p* = 0.0019] (**Figure 1G**).

**Fig 1.**
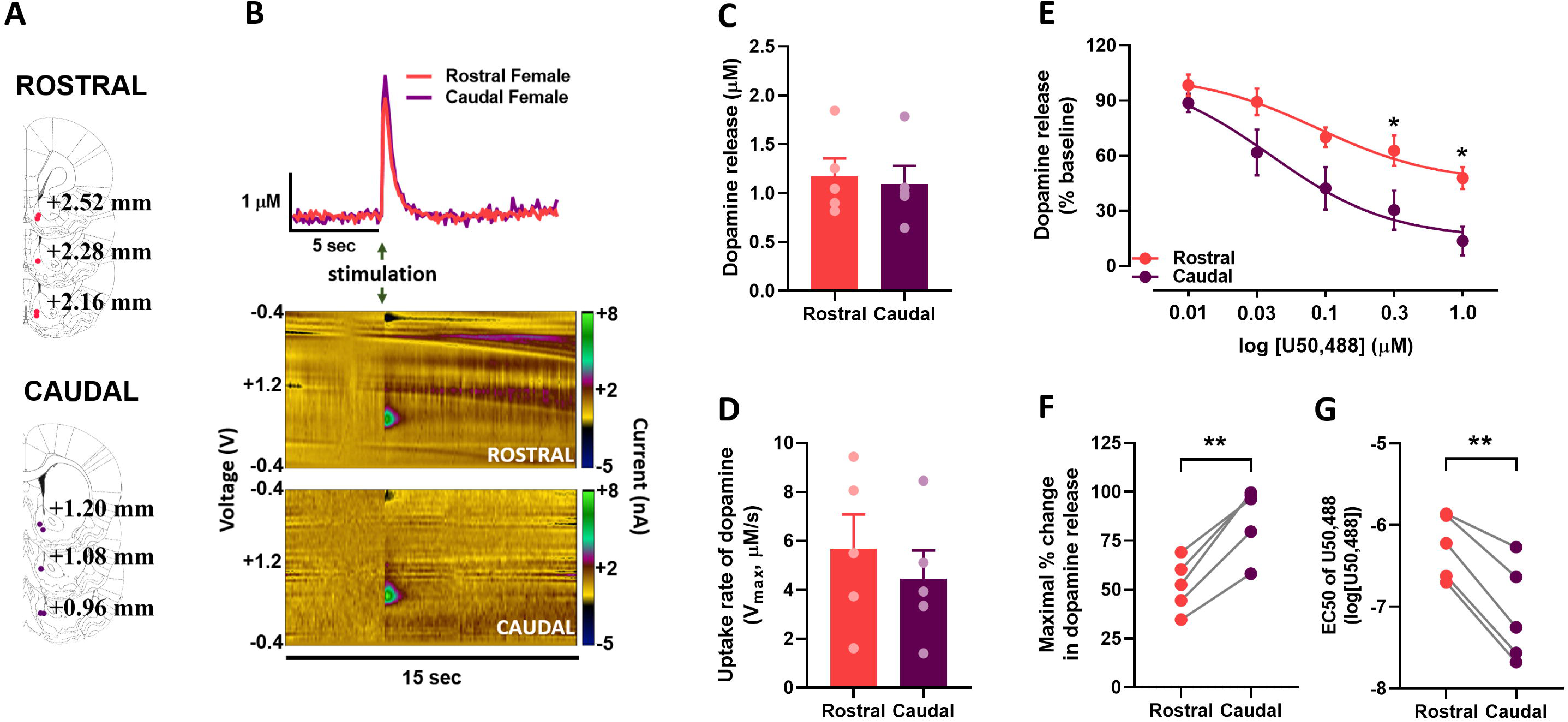
Stimulated dopamine release and impact of KOR activation on dopamine in the rostral and caudal NAc shell in female rats. (A) Carbon-fiber recording electrode placement locations in rostral and caudal subregions of the NAc shell in female rats (*n* = 5; red, rostral; purple, caudal). Because this was a within subject design, one rostral and one caudal slice was used from each rat. (B) Representative traces of transient dopamine signals in response to single pulse electrical stimulation (top; red, rostral; purple, caudal) and their associated colorplots showing transient dopamine signal: *y*-axis, scanning potential; *x*-axis, time; *z*-axis (scale bar), current. (C) There was no significant difference in baseline dopamine release in rostral (red bar) compared to caudal (purple bar) accumbens shell. (D) Dopamine uptake rate was also not different between the rostral and caudal accumbens shell. (E) Function of KOR was significantly greater in the caudal versus rostral accumbens shell. (F) Maximal KOR-mediated reduction in dopamine release was significantly greater in the caudal compared to rostral accumbens shell. (G) Potency of KORs was greater in the caudal compared to rostral accumbens shell. EC50, concentration of drug that results in the half-maximal response in the tissue; KOR, kappa opioid receptor; NAc, nucleus accumbens; U50,488, KOR agonist. ** *p* < 0.01; * *p* < 0.05.

**Fig. 2.**
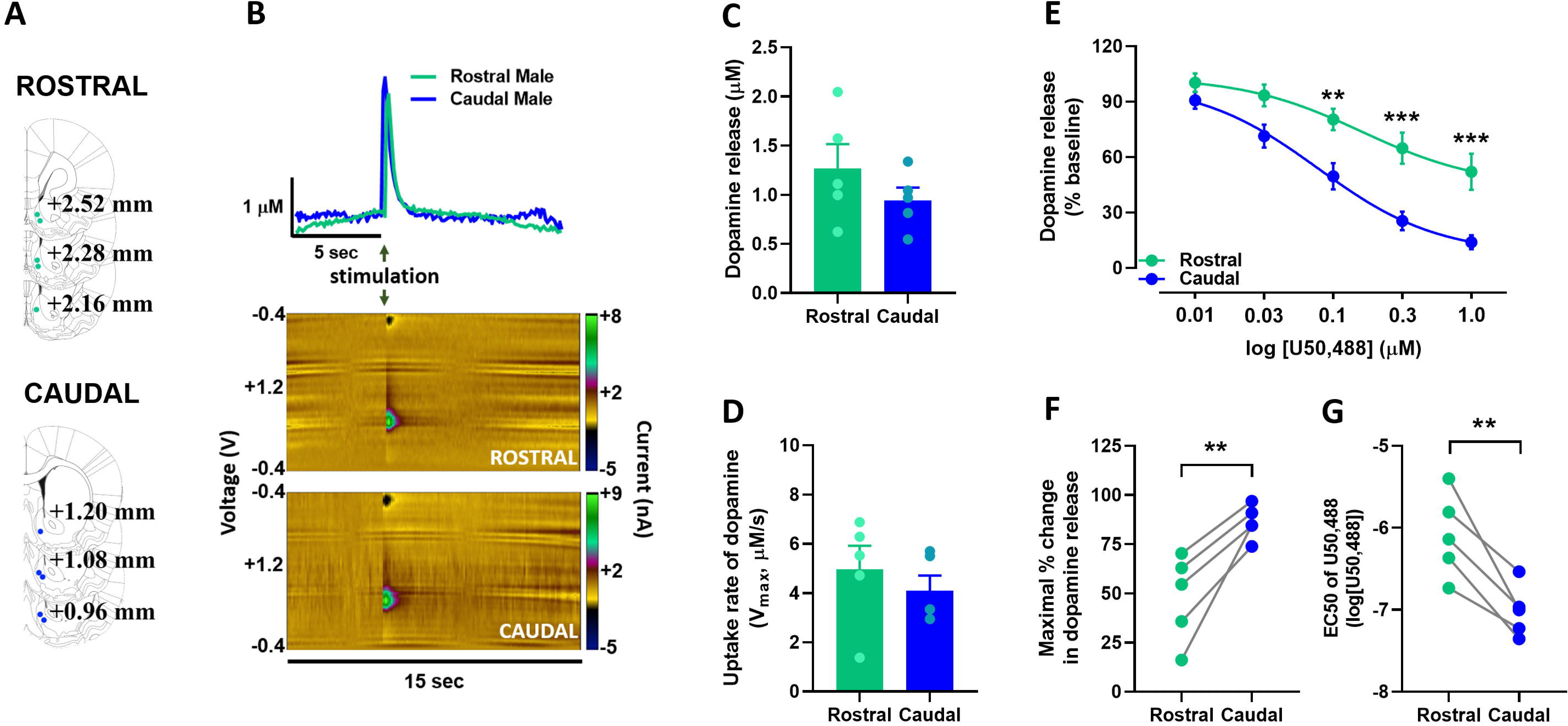
Stimulated dopamine release and impact of KOR activation on dopamine in the rostral and caudal NAc shell in male rats. (A) Carbon-fiber recording electrode placement locations in rostral and caudal subregions of the NAc shell in male rats (*n* = 5; green, rostral; blue, caudal). Because this was a within subject design, one rostral and one caudal slice was used from each rat. (B) Representative traces of transient dopamine signals in response to single pulse electrical stimulation (top; green, rostral; blue, caudal) and their associated colorplots showing transient dopamine signal: *y*-axis, scanning potential; *x*-axis, time; *z*-axis (scale bar), current. (C) There was no significant difference in baseline dopamine release in rostral (green bar) compared to caudal (blue bar) accumbens shell. (D) Dopamine uptake rate did not differ between the rostral and caudal accumbens shell. (E) Function of KOR was greater in the caudal versus rostral accumbens shell. (F) Maximal KOR-mediated reduction in dopamine release was significantly greater in the caudal compared to rostral accumbens shell. (G) Potency of KORs was greater in the caudal compared to rostral accumbens shell. EC50, concentration of drug that results in the half-maximal response in the tissue; KOR, kappa opioid receptor; NAc, nucleus accumbens; U50,488, KOR agonist. *** *p* < 0.001; ** *p* < 0.01.

For experiments in males, electrode placements and representative dopamine signals, along with their respective colorplots, are shown in **Figures 2A** and **2B**, respectively. As in females, comparison of dopamine release between the rostral and caudal NAc shell at baseline (pre-drug exposure) revealed no significant differences [*t*(4) = 1.249; *p* = 0.2798] (**Figure 2C**). Similarly, comparison of dopamine uptake rate (Vmax) between rostral and caudal NAc shell at baseline did not reveal significant differences [*t*(4) = 0.960; *p* = 0.3914] (**Figure 2D**). On the other hand, and similar to females, for KOR-activation mediated inhibition of dopamine release, a repeated-measures two-way ANOVA comparison of rostral and caudal subregions of the NAc shell revealed main effects of U50,488 concentration [*F*(4, 32) = 121.300, *p* < 0.0001] and subregion [*F*(1, 8) = 12.020, *p* = 0.0085], as well as a significant interaction between U50,488 concentration and subregion [*F*(4, 32) = 7.036, *p* < 0.0003]. *Post-hoc* Sidak’s pairwise comparisons revealed that U50,488 at 0.10 µM (*p* = 0.0079), 0.3 µM (*p* = 0.0005), and 1.0 µM (*p* = 0.0007) inhibited dopamine release to a significantly greater extent in the caudal compared to rostral NAc shell (**Figure 2E**). The maximal effect of U50,488-induced inhibition of dopamine release was significantly greater in the caudal compared to rostral subregion of the NAc shell [*t*(4) = 4.875; *p* = 0.0082] (**Figure 2F**). A paired *t*-test comparison of the EC50 between rostral and caudal regions also showed a significant difference [*t*(4) = 4.959, *p* = 0.0077] (**Figure 2G**). Direct comparison between males and females revealed no significant sex differences in dopamine kinetics, KOR function, efficacy, or potency. These data indicate that, in both males and females, while KOR activation inhibits dopamine release in both the rostral and caudal subregions of the NAc shell, the effect is greater in the caudal subregion. Furthermore, this effect appears to be driven by a difference in both efficacy, shown by greater maximal effect in inhibition of dopamine release, and by potency, demonstrated by a reduction in EC50.

### 3.2. Experiment 2: Kappa-opioid stimulation in caudal NAc shell enhances novelty-induced rearing behavior

To determine the effects of KOR stimulation on novelty-induced locomotor activity, rats (*N* = 33) were tested in a novel activity chamber following injection in the rostral or caudal NAc shell with U50,488 or saline vehicle (between-subject) (**Figure 3A**). While unpaired *t*-tests revealed that U50,488 compared to saline had no significant effect on vertical counts when injected in the rostral shell [*t*(19) = -0.575, *p* = 0.5718], it significantly increased them when injected in the caudal shell [*t*(19) = -2.981, *p* = 0.0138] (**Figure 3B**). Examination of ambulatory distance revealed that U50,488 compared to saline again had no significant effect in the rostral shell [*t*(19) = 0.355, *p* = 0.7267] but significantly reduced ambulatory distance when injected in the caudal shell [*t*(19) = 4.255, *p* = 0.0017] (**Figure 3C**). These results demonstrate that U50,488 increases rearing behavior and decreases distance traveled in a novel activity chamber when injected into the caudal but not rostral NAc shell.

**Fig. 3.**
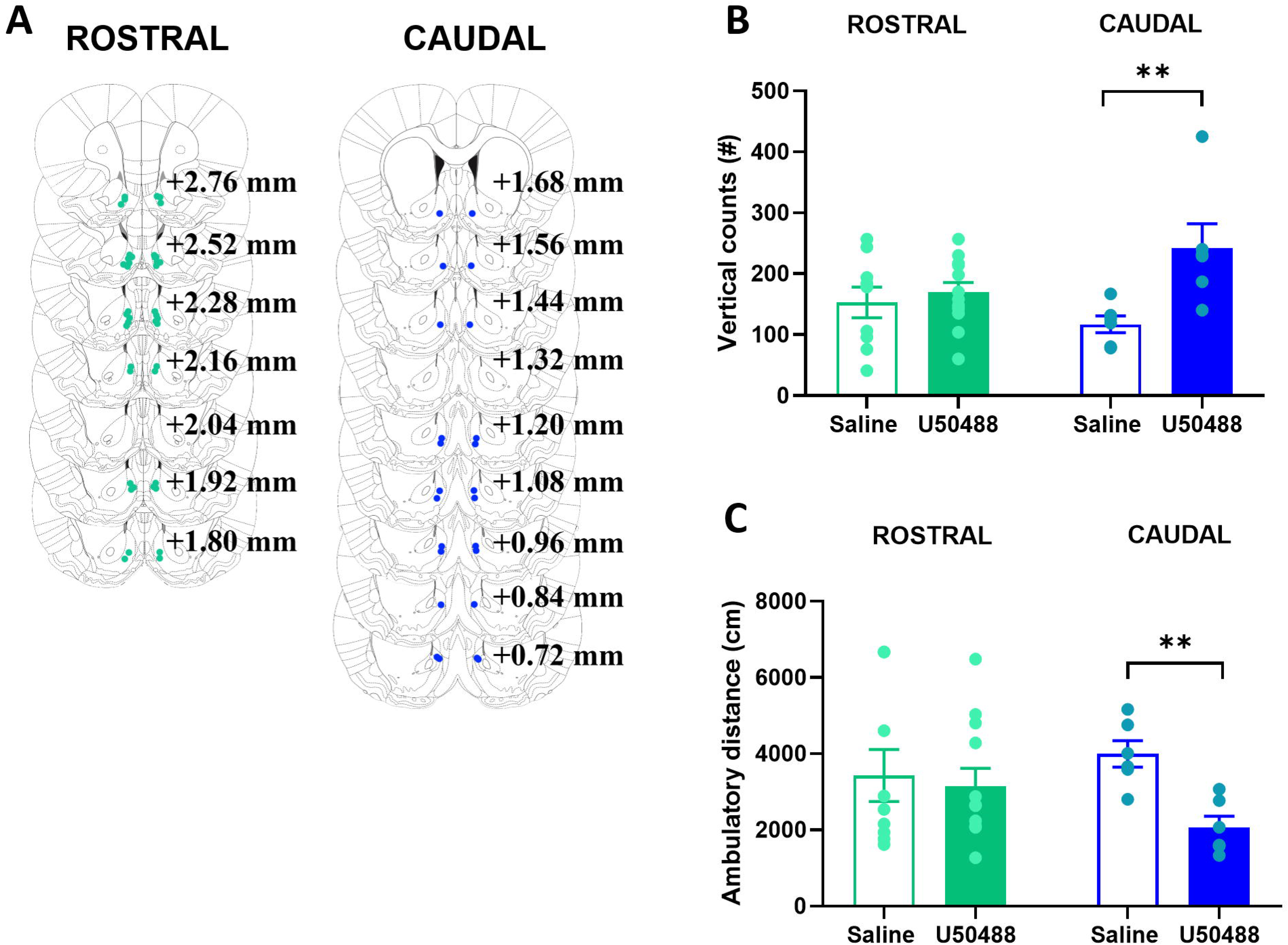
Impact of KOR activation in the rostral and caudal NAc shell on behavior in a novel activity chamber. (A) Microinjection locations for novelty-induced locomotor activity testing (green dots, male rostral; blue dots, male caudal). A total of 21 rats received injections in the rostral NAc shell (*n* = 9 – 12/treatment) and 12 received them in the caudal NAc shell (*n* = 6/treatment). (B) Stimulation of KORs in the caudal but not rostral accumbens shell increased rearing behavior in a novel activity chamber, measured as vertical counts. (C) Stimulation of KORs in the caudal but not rostral accumbens shell reduced distance traveled in a novel activity chamber, measured as ambulatory distance. KOR, kappa opioid receptor; NAc, nucleus accumbens; U50,488, KOR agonist. ** *p* < 0.01.

### 3.3. Experiment 3: Kappa-opioid stimulation in caudal NAc shell induces avoidance behavior in a light-dark box

To determine the effects of KOR stimulation on approach-avoidance behavior, rats (*N* = 17) were tested in a light-dark box following injection in the rostral or caudal NAc shell with U50,488 or saline vehicle (within-subject) (**Figure 4A**). Paired *t*-tests on time spent in the light chamber revealed that U50,488 compared to saline led to a non-significant increase following injection in the rostral shell (+17%, [*t*(10) = -1.551, *p* = 0.1519]) and a significant reduction following injection in the caudal shell [*t*(5) = 2.926, *p* = 0.0328] (**Figure 4B**). Examination of number of entries into the light chamber revealed no significant effects with injection in either the rostral shell [*t*(10) = 1.053, *p* = 0.3170] or the caudal shell [*t*(5) = 1.101, *p* = 0.3210] (**Figure 4C and 4D**). These results suggest that U50,488 induces avoidance behavior in a light-dark box when injected into the caudal but not rostral NAc shell.

**Fig. 4.**
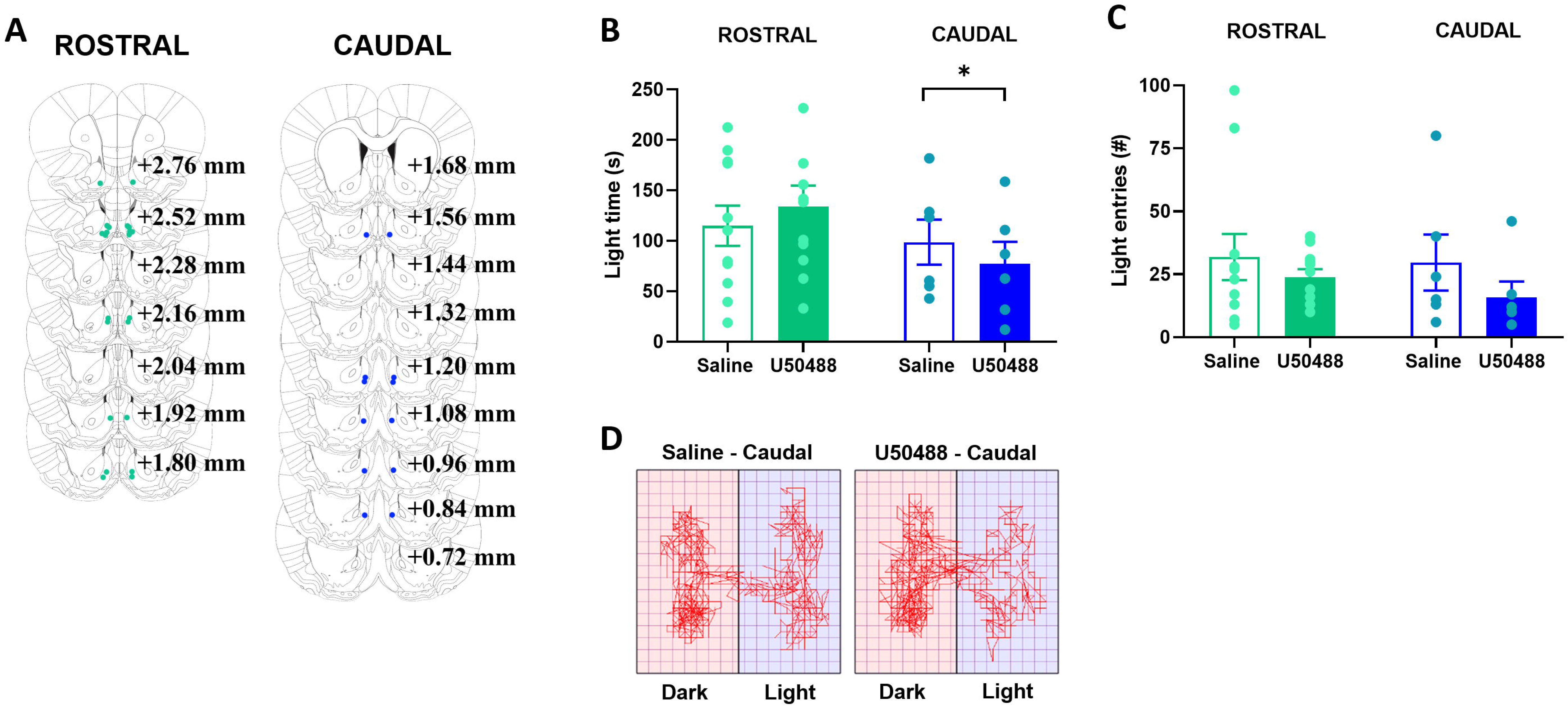
Impact of KOR activation in the rostral and caudal NAc shell on behavior in a light-dark box. (A) Microinjection locations for light-dark box testing (green dots, male rostral; blue dots, male caudal). Rats were injected in the rostral NAc shell (*n* = 11) or caudal NAc shell (*n* = 6). (B) Stimulation of KORs in the caudal but not rostral accumbens shell reduced time spent in the light chamber of a light-dark box. (C) Stimulation of KORs in neither the rostral nor the caudal accumbens shell affected the number of entries into the light chamber of a light-dark box. (D) Representative activity traces of a rat injected in the caudal accumbens shell with saline (left) or U50,488 (right). KOR, kappa opioid receptor; NAc, nucleus accumbens; U50,488, KOR agonist. * *p* < 0.05.

### 3.4. Experiment 4: Kappa-opioid stimulation in rostral NAc shell induces approach behavior in an open field

To determine the effects of KOR stimulation on another measure of approach-avoidance behavior, rats (*N* = 20) were tested in an open field following injection in the rostral or caudal NAc shell with U50,488 or saline vehicle (within-subject) (**Figure 5A**). Paired *t*-tests on time spent in the center area revealed that U50,488 compared to saline had no significant effects when injected in either the rostral shell [*t*(11) = 0.735, *p* = 0.4775] or the caudal shell [*t*(6) = 1.118, *p* = 0.3065] (**Figure 5B**). Examination of number of entries into the center, however, revealed that U50,488 compared to saline led to a significant increase in the number of center entries following injection in the rostral shell [*t*(11) = -3.347, *p* = 0.0065] and to a non-significant decrease when injected in the caudal shell (−24%, [*t*(6) = 0.560, *p* = 0.5957]) (**Figure 5C**). Examination of ambulatory distance revealed that U50,488 compared to saline non-significantly increased ambulatory distance following injection in the rostral shell (+18%, [*t*(11) = -1.399, *p* = 0.1894]) and significantly reduced it following injection in the caudal shell [*t*(6) = 4.222, *p* = 0.0056] (**Figure 5D and 5E**). These results suggest that U50,488 induces approach behavior in an open field when injected into the rostral shell and reduces locomotor activity when injected into the caudal shell.

**Fig. 5.**
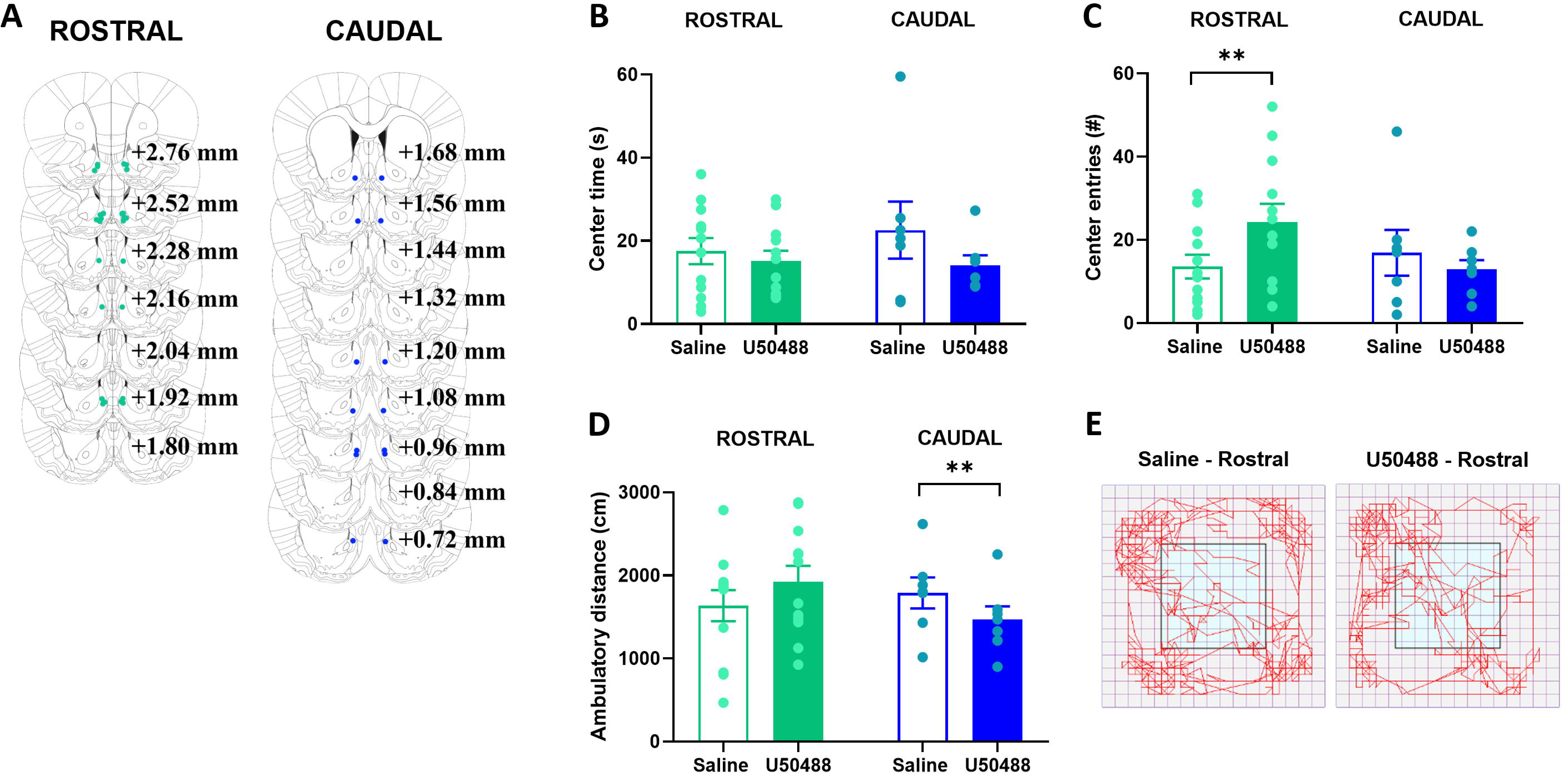
Impact of KOR activation in the rostral and caudal NAc shell on behavior in an open field. (A) Microinjection locations for open field testing (green dots, male rostral; blue dots, male caudal). Rats were injected in the rostral NAc shell (*n* = 13) or caudal NAc shell (*n* = 7). (B) Stimulation of KORs in neither the rostral nor the caudal accumbens shell affected the time spent in the center of an open field. (C) Stimulation of KORs in the rostral but not caudal accumbens shell increased the number of entries into the center of an open field. (D) Stimulation of KORs in the caudal but not rostral accumbens shell reduced distance traveled in an open field. (E) Representative activity traces of a rat injected in the rostral accumbens shell with saline (left) or U50,488 (right). KOR, kappa opioid receptor; NAc, nucleus accumbens; U50,488, KOR agonist. ** *p* < 0.01.

## 4. Discussion

Here, we show that activation of KORs across the rostro-caudal axis of the NAc shell results in different and sometimes opposite effects on dopamine release and behavior. While KOR activation throughout the shell inhibits dopamine release, it does so to a significantly greater extent in the caudal compared to the rostral subregion. Moreover, activation of KORs in the caudal shell increases rearing and avoidance behavior, and reduces locomotor activity, while activation in the rostral shell instead induces approach behavior. These data suggest that KORs in the NAc shell induce different affective behaviors in exploration-based tasks depending on the specific location of their activation, possibly due to their differential level of dopamine inhibition.

While KOR activation with U50,488 inhibited electrically stimulated dopamine release across the NAc shell, this effect was greater in the caudal compared to the rostral shell. This regionally differential effect appears to be a result of increased potency as well as efficacy in the caudal shell. As dopamine transmission itself was equivalent in the two subregions, as measured by dopamine kinetics, it appears unlikely that the KOR-induced differences in dopamine release were due to different degrees of dopamine innervation. Indeed, tract tracing studies have shown that the mesolimbic projection is equally dense across the subregions, although the projection neurons themselves may be genetically distinct [34]. It is unlikely that the KOR receptor is structurally different between the subregions; nevertheless, the affinity of the agonist to the receptor appears to be greater in the caudal shell, as shown by a reduction in EC50 of U50,488. Earlier work with the neuropeptide, cholecystokinin, has similarly found subregion-specific effects on levels of dopamine, despite the presumed presence of cholecystokinin receptors in both the rostral and caudal NAc [35]. Ligand affinity can be influenced by many factors, including cellular milieu and properties of the lipid bilayer, which alter G-protein coupling [36,37]. Therefore, the intracellular cascade triggered by U50,488 binding may itself be different along the rostro-caudal axis of the NAc shell.

In the caudal NAc shell, KOR activation not only significantly reduced dopamine release, but it also increased rearing behavior in a novel activity chamber and avoidance behavior in a light-dark box, and reduced locomotor activity in both a novel chamber and familiar open field. Increased rearing in a novel environment has been associated with reduced levels of dopamine in the striatum [38] and predicts subsequent high-level ethanol drinking [27]. We postulate that novelty-induced rearing may represent risk-taking or impulsivity. The KOR-stimulated increase in avoidance behavior is consistent with prior observations of its aversive effects in the caudal NAc shell [8] and has previously been associated with reduced dopamine release and activity in the NAc shell [39,40]. Finally, while locomotor activity is commonly thought to involve the NAc core rather than shell, a reduction of dopamine with application of 7-OH-DPAT in the NAc shell has been shown to decrease locomotor activity [41]. Thus, the KOR-induced reduction of dopamine release in the caudal NAc shell could be responsible for the array of behavioral changes observed in the present study.

In the rostral NAc shell, while KOR activation reduced dopamine release to a lesser extent than in the caudal shell, it in fact enhanced approach behavior, as measured by a slight increase in time in the light chamber of a light-dark box and a significant increase in the number of center entries in an open field. These effects of U50,488 in the rostral shell are consistent with prior observations of its positive hedonic effects [8]. The findings of subregion-specific KOR-induced differences in *degree* for dopamine release but *direction* for behavior may parallel effects of corticotropin-releasing factor, where, for example, central injections of this neuropeptide at high doses induce taste aversion but at low doses induce preference [42]. Therefore, the diminished inhibition of dopamine release in the rostral shell could be responsible for promoting positive affect.

An alternative explanation for the opposing behavioral effects observed in the current study is that KOR activation modulates several neurochemical systems within the NAc shell that together affect the behavioral outcomes. The NAc receives glutamatergic input from the amygdala, prefrontal cortex, and hippocampus [43,44], and KOR-mediated selective inhibition of amygdalar glutamate transmission in the NAc attenuates the excitatory drive on local D1 dopamine receptor-containing medium spiny neurons [45]. In addition, strong cortical activation-mediated suppression of hippocampal terminal activity in the NAc is reduced by KOR blockade [46], which could impact the D2 dopamine receptor-containing medium spiny neurons [44]. Therefore, KOR activation could differentially affect dopaminergic and glutamatergic multi-synaptic afferents into the rostral compared to caudal NAc shell and this, in turn, could differentially affect efferent signaling.

## 5. Conclusions

Together, our results indicate that KOR stimulation across the rostro-caudal axis of the NAc shell results in different levels of inhibition of dopamine release and either avoidance or approach behavior. While direct tests remain to be conducted to demonstrate that these opposing changes in behavior are *caused* by changes in dopamine dynamics, the data nevertheless demonstrate that the anatomical heterogeneity of the NAc shell extends to differences in the effects of KOR stimulation on dopamine, and in exploration-based tasks as well as affective behavior.

## Acknowledgements

We thank Zachary R. Merkle (Drexel University) for his excellent technical assistance and Dr. Michael Bruchas (University of Washington) for his input on this project.

## Author contributions

**Breanne E. Pirino:** Conceptualization, Formal analysis, Investigation, Writing – review & editing. **Mary B. Spodnick:** Conceptualization, Formal analysis, Investigation, Writing – review & editing. **Andrew T. Gargiulo:** Conceptualization, Investigation, Writing – review & editing. **Genevieve R. Curtis:** Conceptualization, Investigation, Writing – review & editing. **Jessica R. Barson:** Conceptualization, Formal analysis, Funding acquisition, Project administration, Visualization, Writing – original draft, Writing – review & editing. **Anushree N. Karkhanis:** Conceptualization, Formal analysis, Funding acquisition, Project administration, Visualization, Writing – original draft, Writing – review & editing.

KOR: kappa opioid receptor
NAc: nucleus accumbens

